# Indicators of a data-deficient taxa: combining bird and environmental data enhances predictive accuracy of wild bee richness

**DOI:** 10.1101/2024.02.14.580016

**Authors:** Josée S. Rousseau, Alison Johnston, Amanda D. Rodewald

## Abstract

Widespread declines in wild bee populations necessitate urgent action, but there remains insufficient data to guide conservation efforts. Addressing this data deficit, we investigated the relative performance of environmental and/or taxon-based indicators to predict wild bee richness in the eastern and central U.S. Our methodology leveraged publicly available data on bees (SCAN and GBIF data repository), birds (eBird participatory science project) and land cover data (USGS Cropland Data Layer). We used a Bayesian variable selection algorithm to select variables that best predicted bee richness using two datasets: a semi-structured dataset covering a wide geographical and temporal range and a structured dataset covering a focused extent with a standardized protocol. We demonstrate that an indicator based on the combination of bird and land cover data was better at predicting wild bee richness across broad geographies than indicators based on land cover or birds alone, particularly for the semi-structured dataset. In the case of wild bees specifically, we suggest that bird and land cover data serve as useful indicators to guide monitoring and conservation priorities until the quality and quantity of bee data improve.

## INTRODUCTION

There is an urgent need to protect populations of wild bees, as concerns grow about their decline and the risk of losing the ecological roles and ecosystem services they provide (Bartomeus et al., 2013; Cameron and Sadd, 2020; Grixti et al., 2009; IPBES, 2016; Zattara and Aizen, 2021). Unfortunately, conservation action continues to be stymied by the paucity of rigorous information on bee populations and communities (Rousseau et al., 2023; Winfree, 2010). Despite several initiatives to fill data gaps (e.g., Droege et al., 2016; Woodard et al., 2020) and increased numbers of observations submitted to participatory science projects like iNaturalist, wild bees are likely to remain data deficient in the near term. When facing such information needs, a common approach has been to develop indicators that can be used to understand populations or communities (Chase et al., 2000; Fleishman et al., 2005), evaluate environmental conditions (Bryce et al., 2002; Burger, 2006; Hilty and Merenlender, 2000; Niemi and McDonald, 2004), and/or inform management (Pérez-Fuertes et al., 2016; Petrou and Petrou, 2011; Terrigeol et al., 2022). Indeed, previous research demonstrates the usefulness of indicators based on a wide range of taxa, including mammals (Chase et al., 2000; Tognelli, 2005; Yong et al., 2016), butterflies (Fleishman et al., 2005; Rossi and Van Halder, 2010), fish (Roset et al., 2007), and birds (Basile et al., 2021; Chase et al., 2000; Drever et al., 2008). However, less is known about the relative effectiveness of different approaches to developing indicators and, specifically, whether they can be reliably built with environmental or taxon-based variables (Carmel and Stoller-Cavari, 2006; Mandelik et al., 2012).

Environmental indicators or surrogates of biodiversity can include metrics describing ecosystems or landscapes, such as vegetation indices (e.g., NDVI), habitat heterogeneity, structural complexity, land use cover, or topography (Heink and Kowarik, 2010; Niemi and McDonald, 2004; Sowińska-Świerkosz, 2020). The underlying rationale for using environmental surrogates is sound, given that aspects such as land cover classes, can reflect habitat or landscape conditions that affect species. One clear advantage of environmental surrogates is the ease with which one can access a variety of remotely sensed data representing broad spatial extents and different time periods (e.g., including infrared; Nagendra, 2001; Rocchini et al., 2015). However, the resolution and detail of remotely-sensed data are often coarse and insufficient to describe ecological attributes required by any given species. In particular, satellite imagery is unlikely to capture microhabitat features, species interactions (e.g., the presence of competitors or predators), or land management practices (Galbraith et al., 2015; Rocchini et al., 2015). These limitations might be resolved, in part, by using data on other species that may capture multiple dimensions of habitat as well as species interactions better than environmental surrogates (Fleishman et al., 2018; Rodrigues and Brooks, 2007, but see Mandelik et al., 2012).

Taxon-based indicators use data on a single species, an assemblage of species, or an ecological community as proxies to represent other species or indirectly describe aspects of the environment that are difficult to measure directly (Bal et al., 2018; Landres et al., 1988). These indicators are most effective when based upon species that are relatively common, easily detected, and cost-effective to sample (Carignan and Villard, 2002; McGeoch, 1998). Among animal taxa, invertebrates have been used as proxies for environmental health (Siddig et al., 2016) while birds are typically used to assess biodiversity and environmental quality (Fraixedas et al., 2020; Johnson, 2007; Mekonen, 2017). Advantages of using birds is that they are common, easy to survey, strongly associated with habitat and landscape attributes, and are affected by processes operating across multiple scales (Carignan and Villard, 2002; Gardner et al., 2008; Ikin et al., 2014; Niemi et al., 2004). Moreover, the proliferation of participatory science projects like eBird, have made birds unrivaled in terms of data availability over time and space and at low cost (Kosmala et al., 2016; McKinley et al., 2015; Munson et al., 2010;

Theobald et al., 2015). Though single species have been successfully used as proxies (Bustos-Baez and Frid, 2003; De Cáceres et al., 2010; Favreau et al., 2006; Halme et al., 2009), indicators based on species assemblages are generally recommended and considered to perform better (De Cáceres et al., 2012; Dufrêne and Legendre, 1997; Sewell and Griffiths, 2009; Valente et al., 2022), especially when species collectively represent a range of life histories, habitats, and sensitivities to habitat modifications and disturbances (Carignan and Villard, 2002; Fleishman et al., 2018, 2005). These assemblages of individual species can represent multiple taxa, as with Management Indicator Species used by USDA Forest Service (Unkel, 1985; e.g., Moseley et al., 2010). In contrast, indicators built from community metrics, like species richness, have had limited success at characterizing patterns of species of interest (Eglington et al., 2012; Wolters et al., 2006).

Surprisingly few examples exist for indicators that combine environmental and species indicators, despite the potential to leverage the advantages of each. Ferris and Humphrey (1999) alluded to using indicator species in combination with habitat structures as ‘potential indicators of biodiversity’, however, to our knowledge, Fleishman et al. (2018) was first to document that a combination of environmental variables and indicator species best explained variation in species richness.

Here we investigate which combination of environmental and taxon-based data best predicts species richness of wild bees in the eastern and central U.S.. Concern about wild bees continues to rise as populations decline, species are extirpated, and critical habitat resources are lost, yet data deficiencies still limit our ability to detect and respond to changes. The convergence of urgency to act and limited data upon which to base actions makes bees a group for which indicators are likely to be valuable. Previous research used expert-identified and remotely-sensed land cover classes to indicate wild bee abundance across the U.S. (Koh et al., 2016; Lonsdorf et al., 2009). However, many habitat resources used by bees, such flowering plants or ground characteristics (Antoine and Forrest, 2021; Patrício-Roberto and Campos, 2014), are not amenable to detection by satellites (Galbraith et al., 2015). Likewise, a variety of stressors, including pesticides (Janousek et al., 2023; Kennedy et al., 2013; Main et al., 2020) and climate change (Hung et al., 2021; Janousek et al., 2023), may not be evident from remotely-sensed data. Because many bird species are sensitive to multiple spatial scales (e.g., microhabitat, stand, landscape, and region; Frey et al., 2016; Ikin et al, 2014; Saab, 1999) and land management practices (Butler et al., 2010; Jansen and Robertson, 2001), we hypothesized that combining bird and land cover data would best predict the resources, habitats, and landscapes that are associated with diverse bee communities. Here, we compared the performance of indicators of bee richness that were constructed from data on birds, land cover, or a combination of both. Our intention was to develop a tool to guide monitoring, land management, and conservation efforts for bees across large spatial scales until sufficient bee data becomes available.

## METHODOLOGY

### Bee, bird, and land cover data

We used publicly-available and field-based bee and bird data collected in the eastern and central regions of the US (Figure 1) to predict species richness of wild bees. We compared the relative performance of models including predictors that were based on land cover, bird, or bird plus land cover data using bee data from both structured and semi-structured datasets.

**Fig. 1.**
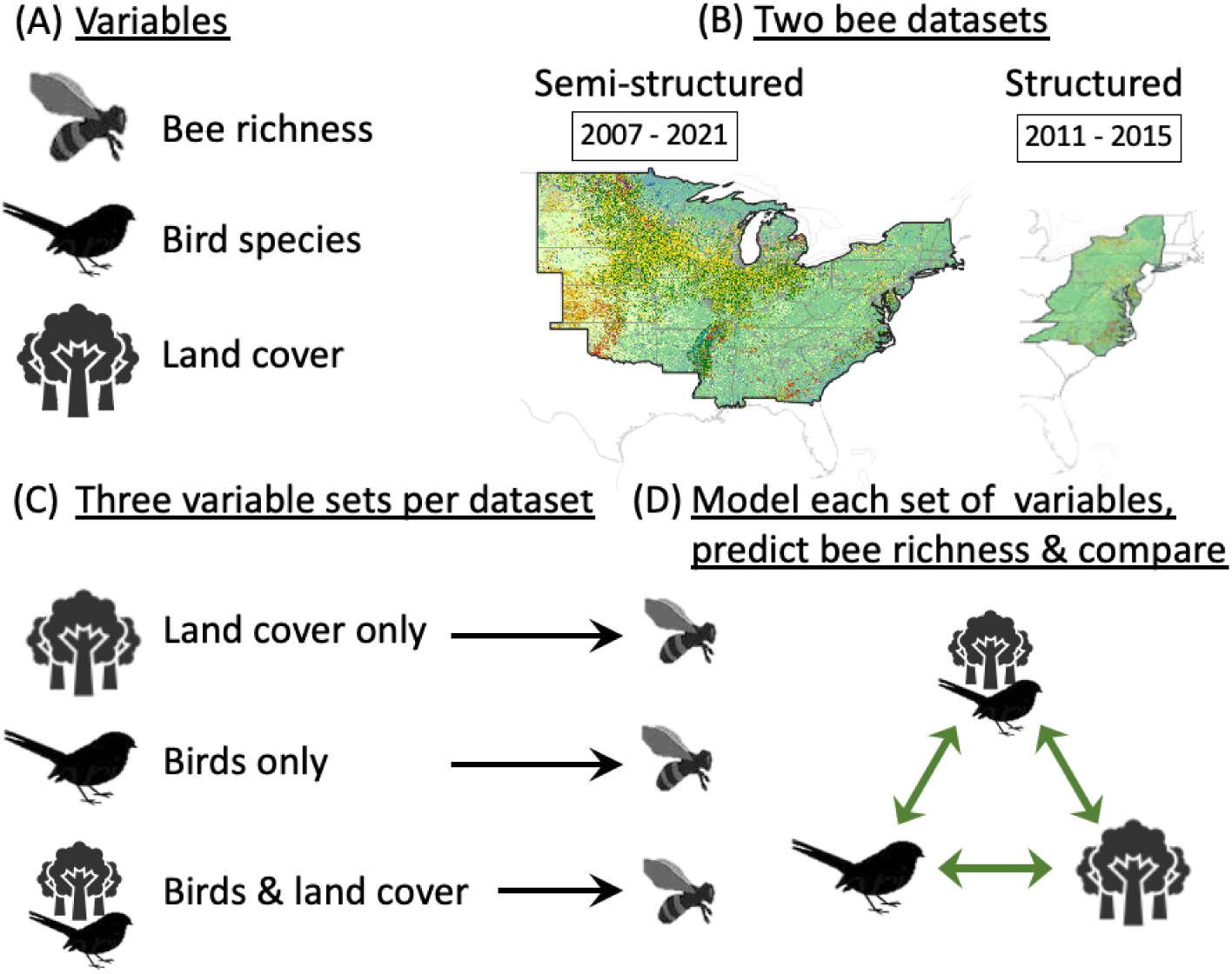
Schema of our methodology. (A) We considered bird species and land cover, to predict bee richness. (B) We used field data from two bee datasets - a semi-structured and structured datasets and summarized them over 3x3 km grid cells, across eastern and midwest U.S.. (C) We compared three sets of variables - land cover only, birds only, and birds and land cover. (D) For each set of variables, we created 100 sub-models using a Bayesian variable selection process. We considered model fit and validation to identify the set of variables that best predicted bee richness. We statistically compared model fit, for each dataset.

The structured dataset consists of data collected using a rigorous protocol and contains information about the bees and associated survey effort (Kelling et al., 2019). It is represented by the U.S. Geological Survey data (Droege and Maffei, 2023), which contains protocol and effort information and could be standardized as the number of bee species per trap in each survey. We selected records of wild bees (excluding honeybees (*Apis mellifera*)) associated with the Bee Inventory and Monitoring Laboratory protocol, which used nets and 3.25 and 12 oz pan traps (Droege et al., 2016). We used surveys where at least 90% of the specimens were identified, and excluded records with missing species identification or with geographic uncertainty exceeding 3 km. We further restricted the temporal range to five years (2011 to 2015) and the geographical extent to a few states in eastern U.S (Figure 1). This produced a dataset with 48,654 bee records, representing 345 species and 1,583 surveys distributed across 390 3x3 km grid cells. We computed the average number of species per survey and trap for each grid cell, as our standardized measure of bee richness.

The semi-structured dataset was sourced from Chesshire et al. (2023), and supplemented with 2021 records from Global Biodiversity Information Facility (GBIF; GBIF.org, 2022) and Symbiota Collections of Arthropods Network (SCAN). Records were collected from 2007 to 2021 in the central and eastern U.S. using a wide range of survey methods and effort The 2021 supplemental data were subject to the same checks, filters, and species name validations as described in Chesshire et al. (2023). We also removed records that were duplicate, lacked species identification, location, or date, or for which uncertainty about geographic location exceeded 3km. This gave us a dataset of 476,584 bee records, representing 792 species across 26,673 3x3 km grid cells. For each grid cell, we calculated the number of species per survey, where a survey was defined by a unique combination of latitude, longitude, and date. Surveys with only one bee, as was the case for most iNaturalist submissions, were excluded as were grid cells with only one survey and fewer than 30 total bee records (Luan et al., 2020; Stockwell and Peterson, 2002; Wisz et al., 2008). For each grid cell, we calculated the mean number of species per survey as a standardized metric of bee richness.

Bird data were extracted from the eBird Basic Dataset (EBD; eBird Basic Dataset, 2022), which consists of bird species checklists submitted by volunteers and subsequently reviewed by experts (Lagoze, 2014; Sullivan et al., 2014). Only records in grid cells associated with bee data in each dataset, were extracted. We selected checklists collected during the bird breeding season (mid-May to mid-August) using stationary or traveling protocols lasting five to 300 minutes. We excluded surveys that did not record counts for all species and those submitted by observers who had submitted fewer than three checklists within our dataset. Bird species abundance data were standardized, within each dataset, accounting for checklist variation in survey effort, time of day, and protocol (supplementary materials S1). We fit a Generalized Additive Model (GAM) for each species separately. The response was the species count per checklist and predictors were: survey duration, survey distance, time the observation started, and protocol. For each checklist and dataset, we calculated species-specific residuals on the log scale from this average relationship. This allowed us to characterize whether checklists recorded high or low species counts, accounting for the checklist effort. For each species and dataset, we then averaged across all checklists within a grid cell, to calculate a mean residual per grid cell, where a positive residual indicates grid cells where a species was more abundant than expected.

To avoid constructing an indicator based on rare species, we established prevalence thresholds to ensure that models included only those bird species that were detected in at least 20% in the grid cells, per dataset (McPherson et al., 2004) and had a breeding distribution covering at least 40% of each study area. The reason for this is that rare species are (by definition) rarely observed and therefore more likely to cause overfitting rather than true associations in our models. One exception to these thresholds were grassland obligate species, for which several were included despite being slightly below the 20% prevalence, because we were especially interested in agricultural landscapes and due to *a priori* ecological expectations. We excluded all records from species that are typically detected as flyovers (e.g., many raptors and some aerial feeders, supplementary material S2) because they could not be linked to local habitat conditions. A total of 79 bird species were considered in our models with the semi-structured dataset and 72 with the structured dataset.

Land cover data were sourced from Cropland Data Layer (CDL), a geo-referenced 30-meters resolution raster, originally obtained from satellite, and categorized into crop-specific land covers (USDA National Agricultural Statistics Service Cropland Data Layer, 2021). We aggregated the CDL from 120 to 45 categories relevant to bee ecology (following Koh et al., 2016) and calculated the percentage of each land cover per 3x3 km grid cell. In our models, we considered only the most common land cover predictors that had a prevalence of at least 20% within each study area. We used data from the 2021 CDL in association with the semi-structured dataset analysis and from 2013 for the structured one.

### Modeling

We modeled bee richness within each 3x3 km grid cell, separately for each dataset. Modeling with the semi-structured dataset incorporated 79 bird species and 21 land cover variables from 2,585 grid cells that contained data on bees, birds, and land covers and met our threshold for analysis. The structured dataset included 72 bird species and 20 land cover variables across 194 grid cells.

After scaling each variable, we used a Bayesian variable selection process from the R package leaps (Miller and Lumley, 2020) to select the best predictors of bee richness using three different sets of candidate predictor variables: (a) land cover only, (b) birds only, or (c) a combination of land cover and birds (Figure 1). For each set of predictors we created 100 sub-models, containing 10 predictors each. We calculated the predicted bee richness within each grid cell by averaging the bee richness predictions from these 100 sub-model predictions. This created three modeled predictions of bee richness, each created with a different set of candidate predictor variables. Sub-models with a large number of potential predictors took a prohibitively long time to run, therefore we used a variable subset selection process to empirically remove predictors that were unlikely to be useful in predicting bee richness (supplementary materials S3). We conducted a sensitivity analysis to ensure that this variable selection process would not impact the results (supplementary materials S3). We also assessed the presence of multicollinearity among predictors in each sub-model using variance inflation factors (VIF). No sub-models had predictors with a Variance Inflation Factor (VIF) of 5 or higher, suggesting minimal multicollinearity (Akinwande et al., 2015).

We assessed the accuracy of our three models separately for each dataset using a five-fold cross-validation process. We re-ran the modeling procedure with each of five subsets of 80% of data, each time creating 100 sub-models and averaging over sub-model predictions to produce modeled predictions for each grid cell within the 20% validation data. This ensured that we were assessing predictions using independent data to prevent positive conclusions being driven by overfitting to the modeled data. We repeated this process five times to create predictions for every grid cell in the original dataset. We repeated the whole procedure for models constructed from each of the three sets of predictor variables: (a) land cover only, (b) birds only, or (c) a combination of land cover and birds.

We compared observed to predicted values of bee richness within each grid cell (plot in supplementary material S4) using a correlation coefficient. We statistically compared the correlation coefficients between observed and predicted bee richness, for models constructed from each of the three sets of predictor variables – land cover only, birds only, birds & land cover. We used the correlation coefficient tests proposed by Hittner et al., (2003) and available through the R package cocor (Diedenhofen and Musch, 2015), to determine which set of variables best predicted observed bee richness, within each dataset.

The best set of variables was used to predict bee richness across the study area associated with each dataset. In order to do this, we needed estimates of relative bird abundance in all grid cells, not only those with eBird checklists, therefore we used the estimated relative abundance per species per grid cell from eBird Status Data Products (Fink et al., 2022). The bee richness point estimates and associated map represent the mean prediction from the 100 sub-models, at each location. For the bee richness uncertainty from the model selection process, we calculated a 90% confidence interval of the bee richness predictions from the 100 sub-models, at each location. Since the bee richness values are relative, the confidence interval range was normalized by the range of point estimates across the entire study extent. This scaled uncertainty reflects the percentage of the variation across models at a given location, relative to the full range of variation across point estimates within all locations.

## RESULTS

The combination of bird and land cover data yielded the most accurate predictions of bee richness using either the semi structured data (Table 1; semi-structured data model fit R^2^ = 0.14, validation R^2^ = 0.14, n = 2585 grid cells) or structured data (model fit R^2^ = 0.28, validation R^2^ = 0.21, n = 194). Plots of observed and predicted values for these analyses are available in the supplementary material (S4).

**Table 1.** Comparison of model fit and associated five-fold validation coefficient of determination among the three sets of variables: land covers only, birds only, and birds & land covers. The model fits are compared among two datasets: data covering the midwest and eastern USA and subset of high quality data covering eastern USA.

| Region & dataset | Model fits | Land cover only | Birds only | Birds & land cover |
| --- | --- | --- | --- | --- |
| Semi-structured dataset | Model fit - R2 | 0.11 | 0.11 | 0.14 |
| Semi-structured dataset | 5-fold validation - R2 | 0.11 | 0.10 | 0.14 |
| Structured dataset | Model fit - R2 | 0.12 | 0.24 | 0.28 |
| Structured dataset | 5-fold validation - R2 | -0.005 | 0.14 | 0.21 |

The inclusion of both land covers and bird species significantly improved correlation coefficients by >15% compared to using either land cover or birds alone. These improvements were significant for the semi-structured dataset, with the model with both land cover and birds being better than land cover only (p < 0.001) and birds only (p < 0.001). For the structured dataset, the model with both land cover and birds was significantly better than the model with land cover only (p = 0.007), but did not show a significant improvement over the model with birds only (p = 0.35). Model fit for birds and land cover was 2x better and significantly improved (p = 0.01) using the structured dataset compared with semi-structured dataset (Table 1).

Focusing on the models with both birds and land cover predictors, the semi-structured and structured datasets had 9 and 5 land cover variables, respectively, and 20 and 26 birds that were selected in at least one of the 100 sub-models (Table 2 and Supplementary material S5).

**Table 2.** List of predictors selected at least once in each of the 100 models per variables set - land covers only, birds only, and birds & land covers - and their associated mean estimate, standard deviation in the estimates, and number of models in which they were selected. Predictor variables are sorted according to frequency of inclusion in “birds & land cover” models.

| Predictor variables | Mean of estimates, SD (# of models) |  |  |
| --- | --- | --- | --- |
|  | Land cover only | Birds only | Birds & land cover |
| Intercept | 4.76, 0.00 (100) | 4.76, 0.00 (100) | 4.76, 0.00 (100) |
| Double crop | 0.51, 0.03 (100) | . | 0.37, 0.01 (100) |
| Urban low-density | -0.79, 0.15 (100) | . | -0.55, 0.05 (100) |
| Barren | 0.48, 0.06 (98) | . | 0.60, 0.02 (100) |
| Deciduous forest | 0.47, 0.09 (63) | . | 0.60, 0.05 (100) |
| Carolina Wren, *Thryothorus ludovicianus* | . | 0.85, 0.08 (100) | 0.64, 0.09 (100) |
| Idle cropland | 0.35, 0.04 (81) | . | 0.31, 0.04 (72) |
| Common Yellowthroat, *Geothlypis trichas* | . | -0.37, 0.04 (68) | -0.36, 0.06 (67) |
| Black-capped Chickadee, *Poecile atricapillus* | . | -0.37, 0.05 (49) | -0.40, 0.04 (60) |
| Blue Jay, *Cyanocitta cristata* | . | -0.46, 0.05 (95) | -0.36, 0.05 (51) |
| Gray Catbird, *Dumetella carolinensis* | . | 0.42, 0.05 (90) | 0.32, 0.04 (45) |
| Ruby-throated Hummingbird, *Archilochus colubris* | . | -0.26, 0.04 (8) | -0.28, 0.02 (41) |
| Alfalfa | -0.39, 0.05 (77) | . | -0.29, 0.02 (37) |
| Open water | -0.39, 0.06 (100) | . | -0.27, 0.02 (24) |
| Scarlet Tanager, *Piranga olivacea* | . | 0.48, 0.08 (90) | 0.29, 0.03 (23) |
| Warbling Vireo, *Vireo gilvus* | . | -0.41, 0.04 (95) | -0.30, 0.03 (20) |
| Dickcissel, *Spiza americana* | . | -0.29, 0.03 (36) | -0.27, 0.02 (18) |
| Orchard Oriole, *Icterus spurius* | . | 0.35, 0.04 (78) | 0.27, 0.02 (14) |
| European Starling, *Sturnus vulgaris* | . | . | 0.25, 0.02 (6) |
| Grass pasture | 0.10, 0.08 (6) | . | 0.24, 0.02 (5) |
| Chipping Sparrow, *Spizella passerina* | . | 0.33, 0.04 (51) | 0.24, 0.01 (3) |
| American Crow, *Corvus brachyrhynchos* | . | 0.26, 0.02 (12) | 0.24, 0.01 (3) |
| Red-bellied Woodpecker, *Melanerpes carolinus* | . | -0.31, 0.06 (2) | -0.34, 0.03 (3) |
| Rose-breasted Grosbeak, *Pheucticus ludovicianus* | . | -0.18 (1) | -0.27, 0.03 (2) |
| Corn | -0.47, 0.14 (58) | . | -0.27 (1) |
| Northern Cardinal, *Cardinalis cardinalis* | . | -0.61, 0.07 (100) | -0.33 (1) |
| White-breasted Nuthatch, *Sitta carolinensis* | . | -0.31, 0.03 (10) | -0.32 (1) |
| Downy Woodpecker, *Dryobates pubescens* | . | -0.31, 0.01 (8) | -0.26 (1) |
| American Redstart, *Setophaga ruticilla* | . | -0.24, 0.02 (7) | -0.29 (1) |
| Yellow-throated Vireo, *Vireo flavifrons* | . | . | -0.26 (1) |
| Mixed forest | -0.39, 0.07 (87) | . | . |
| Urban high-density | -0.38, 0.11 (53) | . | . |
| Coniferous forest | -0.33, 0.09 (45) | . | . |
| Herbaceous wetland | -0.29, 0.07 (36) | . | . |
| Developed open space | 0.27, 0.03 (20) | . | . |
| Urban medium-density | -0.39, 0.14 (20) | . | . |
| Bean | -0.31, 0.12 (18) | . | . |
| Grain | -0.22, 0.04 (18) | . | . |
| Woody wetland | -0.09, 0.08 (7) | . | . |
| Shrubland | 0.09, 0.01 (5) | . | . |
| Grass | 0.01, 0.01 (4) | . | . |
| Orchard | 0.03, 0.01 (4) | . | . |
| Wood Thrush, *Hylocichla mustelina* | . | 0.37, 0.07 (36) | . |
| Cliff Swallow, *Petrochelidon pyrrhonota* | . | 0.26, 0.02 (29) | . |
| American Robin, *Turdus migratorius* | . | -0.37, 0.10 (11) | . |
| Yellow-billed Cuckoo, *Coccyzus americanus* | . | 0.27, 0.02 (11) | . |
| Blue Grosbeak, *Passerina caerulea* | . | 0.29, 0.05 (6) | . |
| Song Sparrow, *Melospiza melodia* | . | -0.28, 0.02 (4) | . |
| American Goldfinch, *Spinus tristis* | . | -0.22 (1) | . |
| Brown Thrasher, *Toxostoma rufum* | . | 0.17 (1) | . |
| House Sparrow, *Passer domesticus* | . | -0.25 (1) | . |

Those land covers and bird variables that were selected in all 100 sub-models typically had a large effect size, based on their mean coefficients. Five variables were selected within all 100 sub-models using the semi-structured dataset - deciduous forest, barren land, double crop, low-density urban, and Carolina Wren - and two with the structured dataset, grain and Gray Catbird (Table 2 and Supplementary material S5). With the exception of low-density urban landscapes, the most selected variables were positively correlated with bee richness (Table 2; supplementary material S5).

In our study area, bee richness was generally higher on the East Coast along the Appalachian Mountains and lower in the Midwest, particularly around Iowa (Figure 2A). Uncertainty in predicted bee richness was lowest around North and South Dakota, Illinois, and along the Atlantic coast, while it was highest near West Virginia, eastern Kentucky, and southern Missouri (Figure 2B).

**Fig. 2.**
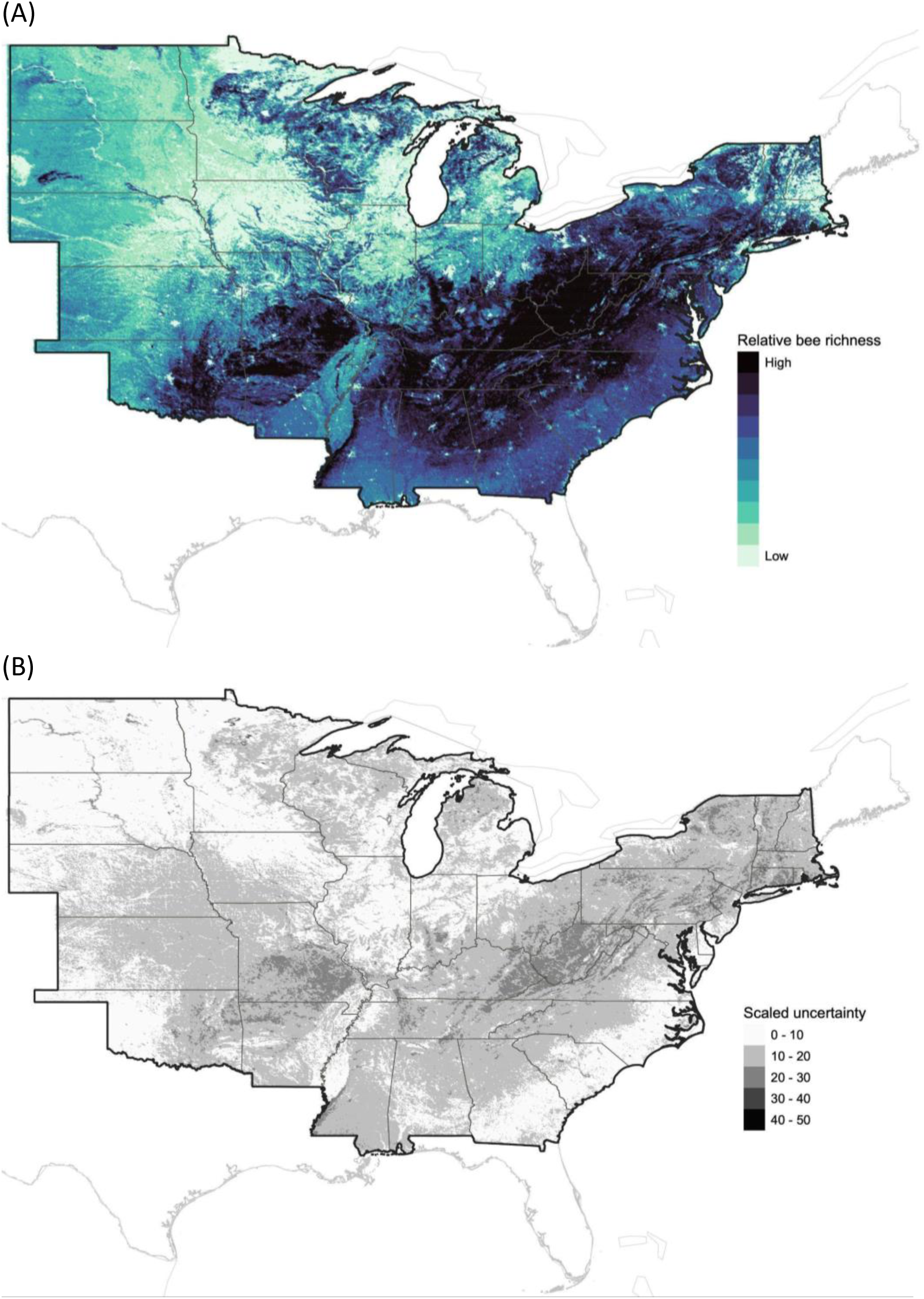
Predicted bee richness using birds and land cover variables. (A) The dark blue represents locations where a relatively higher bee richness is expected, compared to other locations within our study extent. (B) The scaled uncertainty associated with bee richness for each grid cell, where dark color represents higher uncertainty.

## DISCUSSION

Tools for guiding the conservation of data-deficient taxa often include environmental or taxon-based indicators. Though only one type of variable typically is used to create an indicator (but see Fleishman 2017), our results indicate that combining data from both the environment and other taxa may significantly improve the prediction accuracy and, thus, may better inform conservation actions. Unlike previous work that relied upon land cover data to predict bee abundance (Koh et al., 2016; Lonsdorf et al., 2009), we found that bird data added value over land cover alone and improved our ability to predict species richness of wild bees. The usefulness of birds is not surprising, given that they are known to be an effective indicator species for other taxa (Chase et al., 2000; Fleishman et al., 2018, 2005; Rodríguez-Estrella et al., 2019; Thomson et al., 2007).

Several factors may explain why the combination of birds and land cover variables predicted bee richness better than using either land cover or birds alone. First, bird and land cover variables likely offer complementary insights into habitat quality. A broad category of land cover, such as ‘deciduous forest’, usually includes a wide range of floristic composition, habitat structure, patch configuration, age, and land management practices (Milam et al., 2022; Taki et al., 2013; Ulyshen et al., 2023; Urban-Mead et al., 2021). For example, numbers of flowering plants that attract bees are often greater in early-successional than mature forests. Likewise, forests in which understories were replaced by grass (e.g., wooded parks) are unlikely to provide nesting habitat to ground-nesting bees. In such cases, the presence or abundance of particular bird species (e.g., open woodland species like Chipping Sparrow (*Spizella passerina*), forest-understory species like American Redstart (*Setophaga ruticilla*) or Wood Thrush (*Hylocichla mustelina*)) can provide additional information to better identify habitats favored by bees. Indeed, the presence of fruit-eating bird species, such as Gray Catbird (*Dumetella carolinensis*), as predictors highlight the importance of forests containing flowering shrubs and trees for bee communities (Inari et al., 2012; Ramalho, 2004). Second, individual land cover variables may be blind to habitat juxtaposition or the co-occurrence of different habitats in close proximity. For example, while ‘double crops’ may provide sufficient flower resources, bees also may require easy access to less disturbed habitat for nesting (e.g. ‘idle cropland’, ‘grass pastures’, or ‘deciduous forests’) that are better indicated by bird species that nest in trees but forage in open habitats. Third, birds and bees select their habitat based on resources available across multiple scales (Diaz-Forero et al., 2013; Hatfield and LeBuhn, 2007; Orians and Wittenberger, 1991; Pardee and Philpott, 2014; Rollin et al., 2019; Thompson and Mcgarigal, 2002). As such, birds are likely to incorporate multi-scale information relevant to bee population that would not be available through land covers alone. For instance, the presence of species like Orchard Oriole (*Icterus spurius*) can signal the availability of open woodlands, orchards, woody hedgerows, and flowering plants used for nesting and foraging resources by orioles and wild bees alike. Fourth, the combined bird and land cover variables selected as predictors represent a wide range of habitats, which suggest that a higher bee richness may be associated with heterogeneous landscapes (Andersson et al., 2013; Mallinger et al., 2016; Montagnana et al., 2021) that include multiple types of nesting substrates to accommodate ground and cavity nesters and a diversity of flower resources, from crops, shrubs, and trees.

The geographic pattern of wild bee richness predicted by our indicators (Figure 2A) is consistent with previous reports that wild bee richness and/or abundance is highest in landscapes characterized by a mosaic of deciduous forest and low-intensity agriculture and lowest in areas dominated by intensive agriculture, such as the Midwest (Figure 2; Kennedy et al., 2013; Koh et al., 2016). Consistent with the results from Koh et al., (2016), we found high levels of uncertainty in our predictions in certain regions, particularly in areas like the Appalachian and Ozark Mountains (Figure 2B). Given that many bee species require forest habitats during particular life stages (Hanula et al., 2016; Roberts et al., 2017; Smith et al., 2021; Urban-Mead et al., 2021), we were not surprised to find large positive effects of deciduous forests and Carolina Wrens, a species associated with gaps in deciduous forest (Haggerty and Morton, 2020). Also unsurprising were the negative associations we detected between bee richness and intensive agricultural crops, such as corn and alfalfa. Corn monocultures are known to have low bee richness (Gay et al., 2024), in part because of intensive management practices like tilling and pesticide applications, whereas alfalfa is used by relatively few bee genera (Rollin et al., 2013). Importantly, we recognize that species richness does not necessarily indicate conservation value. High species richness could result from communities comprised mainly of generalists and common species, whereas areas of low richness may be home to specialized and rare species might warrant more conservation attention (Bogusch et al., 2020; Raiol et al., 2021; Rousseau et al., 2023; Winfree, 2010). For these reasons, establishing conservation priorities is best done in consultation with experts or, ideally, after ground-truthing with field surveys.

Using one taxon as an indicator for another requires considering ecological context, such as threats affecting both groups, species interactions, and the spatial and temporal scales at which they utilize their habitat. In our case, the breeding territory size and season of most birds align well with the timing and habitat size requirements of many bees. That said, we recognize that bees may require unique resources., For instance, ground-nesting bees may exhibit preferences for specific below-ground resources (Antoine and Forrest, 2021) that may not be well indicated by birds. Additionally, bees are likely influenced by micro-habitats at a finer scale than birds, such as the availability of small bare ground patches. Lastly, the breeding season of birds may include different density-dependent processes compared to bees, where a higher abundance of birds is not always correlated with higher habitat quality (Johnson, 2007). While the association of certain bird species with bee richness may be intuitive, including all bird species *a priori* in our analysis provided insights on novel relationships between these birds, bee richness, and the habitat they occupy.

Our findings also provide insight into the influence of structured versus semi-structured data on results. The improved predictions we generated using the structured dataset are likely due to differences in data quality and scale compared to the semi-structured dataset. The structured dataset included protocol and effort information, enabling us to generate more precise bee richness estimates across space (Johnston et al., 2021; van Strien et al., 2013). Additionally, the use of a limited number of years in the structured dataset minimizes variation in bee and bird species detection due to temporal changes in climate or land cover. The comparatively narrow geographic scope of the structured dataset likely resulted in more consistent species-habitat associations across the study area and, consequently, improved model fits (Rollinson et al., 2021; Rousseau and Betts, 2022). Focusing on a smaller geographical area also increased the likelihood of more bird species having their breeding distribution covering larger portions of the study area. This may be a reason the birds-only model performed relatively better using the structured than semi-structured dataset. Lastly, the model fit using the semi-structured dataset may have been lower because the sample size was much larger and represented a more extensive area in which several regions lacked bee data.

## Conclusion

Recent drastic declines in insect biodiversity (Butchart et al., 2010; Montgomery et al., 2020; Wagner, 2020), underscore a need to use all available information to conserve data-deficient taxa. Despite increases in data availability from sources like satellites or participatory science projects, few have investigated the extent to which integrating data sources may improve the usefulness of indicators of taxonomic groups with limited data. We demonstrated that by combining multiple tools, we can achieve better predictions of bees, which are a data-deficient taxa, but also provide vital ecosystem services. Until more bee data becomes available, our results could be used to guide monitoring efforts, improve conservation of bees through land conservation, and recommend land management practices known to promote healthy bee populations.

## ACKNOWLEDGMENTS

We thank P. Chesshire for sharing information about the quality control and filters that we used when supplementing her compiled dataset and to the following data owners, which provided at least 5% of the records used in the semi-structured dataset: American Museum of Natural History, iNaturalist, University of Kansas Biodiversity Institute, U.S. Department of Agriculture, and U.S. Geological Survey. Thank you to S. Droege for sharing the U.S. Geological Survey data and providing guidance on extracting records associated with the standardized protocol. We appreciate the generous feedback from B. Danforth, the guidance provided by Matt Strimas-Mackey on accessing and manipulating eBird data, and the valuable input from colleagues in the Center for Avian Population Studies at the Cornell Lab of Ornithology that improved our work. Finally, we are especially grateful to our partners at the Cornell Atkinson Center for Sustainability, particularly Patrick Beary and Gail Phillips, for their collaboration, support, and input throughout the project.

