## Supplementary material for "Indicators of a data-deficient taxa: combining bird and environmental data enhances predictive accuracy of wild bee richness"

**SUPPLEMENTAL MATERIAL**

### S1. Standardizing bird abundance using residuals

Bird abundance within each grid cell was standardized separately for each species and dataset. In the first stage of this process, we wanted to account for variation in effort across checklists. We first modeled bird count as a function of survey duration within a Generalized Additive Model. This function describes how bird counts on checklists varied with duration. For most species, the maximum count is reached for checklists of intermediate duration, beyond which counts tend to decline. Although this seems counterintuitive, it is a consistent pattern in many participatory science datasets that likely reflects different behavior of observers or different types of environments associated with longer duration checklists. We wanted to estimate this maximum in order to determine the survey duration (up to 300 minutes maximum) associated with the greatest counts for each species. Survey durations with the maximum counts ranged from 142 to 300 minutes across all the bird species using the semi-structured dataset and 146 to 300 minutes using the structured dataset.

In the second step, we used a log-link poisson Generalized Additive Model, to model bird counts per checklist as a function of effort variables that are likely to affect the observed counts: survey duration, survey distance, the time of the day when the survey started, and the protocol (stationary or traveling). The response variable in the models was the count of the bird species on each checklist, with the predictors being discretized smooth functions of the effort variables. We then calculated the expected count of the bird on each checklist (i.e. the fitted value from a prediction from the GAM model for each set of predictor variables). Next, we took the residual for each checklist: which would be high if the checklist reported more birds of the species than expected once accounting for the effort, and negative if the checklist reported fewer birds of the species than expected once accounting for the checklist effort. We then took the mean residual for all checklists within each grid cell. Positive values indicated when grid cells had higher numbers of that bird species than the average based on effort, whereas negative numbers indicated lower than average counts  This average residual per grid cell was used as a measure of  bird abundance per species  that was used in subsequent models of bee richness.

### S2. List of bird species excluded from analysis

The following species were excluded from the analysis as they are considered flyover species and could not be linked to the local habitat: American Kestrel, Bald Eagle, Broad-winged Hawk, Chimney Swift, Common Nighthawk, Cooper's Hawk, Red-shouldered Hawk, Red-tailed Hawk, Turkey Vulture, Black Vulture.

### S3. Two-step process to select relevant predictors of bee richness

Because computational time increased non-linearly with the number of variables, it was impractical to use the Bayesian model selection procedure with a high number of predictor variables. To overcome this constraint, we employed a two-step process to first select the predictors used in the models to those most associated with bee richness. In order to ensure that this two-step process did not influence the results, for a subset of the models we directly compared the outcomes of this two-step process with those of running a model incorporating all predictors simultaneously. This assessment was conducted on a subset of 78 of 100 variables in the original dataset. This validation confirmed that the two-step process did not influence the results and so we used this for the main results in this paper.

In the initial step, we considered 78 bird species and land cover predictors. We divided the dataset into three subsets, each containing 26 predictors, and then selected the best three predictors in each subset, employing the same Bayesian variable selection methodology used in the model with all 78 predictors. We repeated this process by randomly shuffling the predictor list 1000 times, generating subsets, and selecting the best three predictors each time (we also experimented with using five predictors). We retained all predictors that were chosen at least once across all subsets. These retained predictors were subsequently utilized in the second step to identify the ten predictors most strongly correlated with bee richness. We calculated the model fit and performed a five-fold validation for this model. The results of this assessment indicate that the two-step process yielded the same predictors, predictor estimates, model fit, and validation R-squared as if we had used all predictors simultaneously. However, this two-step process significantly reduced the time required to extract these results, taking only 5 minutes as opposed to several days for the full model.

**Table S3.1.** Comparison of the number of predictors used, resulting model fit and validation r-squared, predictors selected, and associated estimates per methodology tested.

| **Model / process** | **Whole model** | **Whole model** | **Two-step process** | **Two-step process** |
|---|---|---|---|---|
| **Scaling of predictors** | Not scaled | Scaled | Scaled | Scaled |
| **Total # of predictors considered** | 78 | 78 | 78 | 78 |
| **# grid cells** | 571 | 571 | 571 | 571 |
| **# predictors selected per subset** | . | . | 5 | 3 |
| **# predictors retained in two-step process** | . | . | 51 | 41 |
| **# best predictors selected** | 10 | 10 | 10 | 10 |
| **Model fit ($r^2$)** | 0.265 | 0.265 | 0.265 | 0.265 |
| **Five-fold validation ($r^2$)** | 0.224 | 0.224 | 0.224 | 0.224 |
| **Predictors** | **Predictor estimates and significance level** | | | |
| (Intercept) | 0.2412 | 1.2353 *** | 1.2353 *** | 1.2353 *** |
| AmericanRobin | -0.1488 * | -0.1956 * | -0.1956 * | -0.1956 * |
| BrownThrasher | 0.6347 *** | 0.2482 *** | 0.2482 *** | 0.2482 *** |
| HairyWoodpecker | -0.5625 ** | -0.1884 ** | -0.1884 ** | -0.1884 ** |
| pctBean | 0.0260 * | 0.1680 * | 0.1680 * | 0.1680 * |
| pctDeciForest | 0.0150 *** | 0.2837 *** | 0.2837 *** | 0.2837 *** |
| pctHerbWetland | 0.0160 * | 0.1569 * | 0.1569 * | 0.1569 * |
| pctMixedForest | 0.0687 *** | 0.5343 *** | 0.5343 *** | 0.5343 *** |
| pctWoodyWetland | 0.0338 *** | 0.3773 *** | 0.3773 *** | 0.3773 *** |
| RubyThroatedHummingbird | -0.6614 ** | -0.2251 ** | -0.2251 ** | -0.2251 ** |
| WoodThrush | 0.4635 ** | 0.2297 ** | 0.2297 ** | 0.2297 ** |

**S4. Plots of the observed and predicted bee richness for the large-scale and small-scale analysis.**

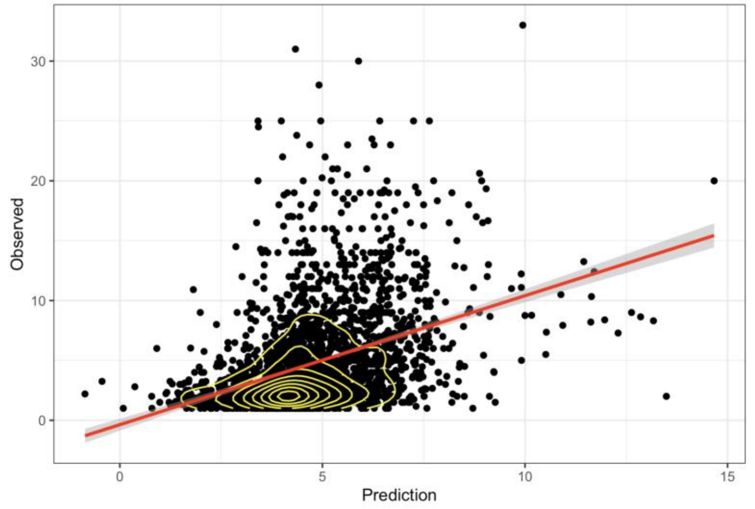


**Fig. S4.1.** Correlation between observed and predicted bee richness for 2,501 locations across the eastern half of the United-State, 2007 to 2021. The yellow contour lines represent the density of points and the red line the linear correlation.

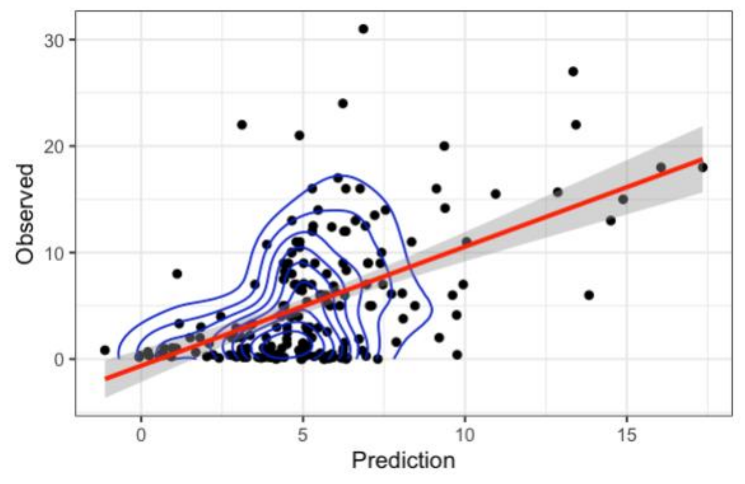


**Fig. S4.2.** Correlation between observed and predicted bee richness for 194 locations across the eastern United-State, 2011 to 2015. The blue contour lines represent the density of points and the red line the linear correlation.

**S5. Structured dataset results**

**Table S5.1.** List of predictors selected at least once in each of the 100 models per variables set - land covers only, birds only, and birds & land covers - and their associated mean estimate, standard deviation in the estimates, and number of models in which they were selected. Predictor variables are sorted according to frequency of inclusion in “birds & land cover” models.

| | **Mean of estimates, SD (# of models)** | | |
|---|---|---|---|
| **Predictor variables** | **Land cover only** | **Birds only** | **Birds & land cover** |
| Intercept | 5.32, 0.00 (100) | 5.32, 0.00 (100) | 5.32, 0.00 (100) |
| Grain | 1.59, 0.09 (100) | . | 1.43, 0.09 (100) |
| Gray Catbird, *Dumetella carolinensis* | . | 1.82, 0.20 (100) | 1.34, 0.19 (100) |
| Yellow-throated Vireo, *Vireo flavifrons* | . | -0.96, 0.10 (99) | -0.98, 0.11 (97) |
| Green Heron, *Butorides virescens* | . | -0.92, 0.12 (72) | -0.92, 0.09 (93) |
| Brown Thrasher, *Toxostoma rufum* | . | -0.96, 0.11 (89) | -0.96, 0.14 (89) |
| Mixed forest | -0.95, 0.03 (100) | . | -0.81, 0.09 (84) |
| Chipping Sparrow, *Spizella passerina* | . | 0.91, 0.13 (85) | 0.85, 0.11 (76) |
| Brown-headed Cowbird, *Molothrus ater* | . | 0.64, 0.09 (14) | 0.76, 0.09 (58) |
| Fish Crow, *Corvus ossifragus* | . | -0.97, 0.13 (85) | -0.83, 0.14 (50) |
| Red-winged Blackbird, *Agelaius phoeniceus* | . | -1.24, 0.24 (65) | -1.02, 0.22 (43) |
| Purple Martin, *Progne subis* | . | -0.84, 0.14 (38) | -0.77, 0.11 (38) |
| Barn Swallow, *Hirundo rustica* | . | 1.25, 0.21 (86) | 1.09, 0.23 (35) |
| House Wren, *Troglodytes aedon* | . | -0.99, 0.17 (73) | -0.74, 0.08 (33) |
| Blue Jay, *Cyanocitta cristata* | . | -0.80, 0.09 (43) | -0.64, 0.07 (18) |
| Eastern Wood-Pewee, *Contopus virens* | . | -0.44 (1) | -0.73, 0.15 (16) |
| Great Blue Heron, *Ardea herodias* | . | -0.39 (1) | -0.63, 0.11 (15) |

| | | | |
|---|---|---|---|
| Coniferous forest | -0.15, 0.04 (10) | . | -0.70, 0.03 (13) |
| Eastern Bluebird, *Sialia sialis* | . | -0.66, 0.08 (13) | -0.65, 0.14 (11) |
| House Finch, *Haemorhous mexicanus* | . | -0.61, 0.12 (6) | -0.57, 0.09 (8) |
| Red-bellied Woodpecker, *Melanerpes carolinus* | . | . | -0.64, 0.04 (6) |
| Herbaceous wetland | -0.51, 0.04 (100) | . | -0.54, 0.08 (3) |
| Alfalfa | -0.33, 0.02 (65) | . | -0.64, 0.06 (3) |
| Northern Rough-winged Swallow, *Stelgidopteryx serripennis* | . | 0.70, 0.11 (30) | 0.48, 0.03 (2) |
| Summer Tanager, *Piranga rubra* | . | . | 0.69, 0.33 (2) |
| Indigo Bunting, *Passerina cyanea* | . | 0.62, 0.21 (7) | 0.47 (1) |
| Common Yellowthroat, *Geothlypis trichas* | . | -0.69, 0.33 (2) | -0.57 (1) |
| Wood Thrush, *Hylocichla mustelina* | . | 0.42 (1) | 1.17 (1) |
| Ovenbird, *Seiurus aurocapilla* | . | -0.28 (1) | -1.31 (1) |
| Orchard Oriole, *Icterus spurius* | . | . | -0.49 (1) |
| Tufted Titmouse, *Baeolophus bicolor* | . | . | -0.51 (1) |
| Pine Warbler, *Setophaga pinus* | . | . | -0.79 (1) |
| Double crop | -0.48, 0.10 (98) | . | . |
| Urban medium-density | 0.48, 0.10 (91) | . | . |
| Woody wetland | -0.42, 0.08 (91) | . | . |
| Shrubland | 0.36, 0.04 (80) | . | . |
| Open water | -0.30, 0.03 (79) | . | . |
| Grass | -0.23, 0.02 (43) | . | . |
| Deciduous forest | 0.27, 0.05 (32) | . | . |
| Corn | 0.31, 0.06 (30) | . | . |

| | | | |
|---|---|---|---|
| Barren | -0.16, 0.02 (18) | . | . |
| Developed open space | -0.20, 0.03 (16) | . | . |
| Urban high-density | -0.13, 0.22 (14) | . | . |
| Urban low-density | -0.04, 0.25 (11) | . | . |
| Bean | -0.17, 0.09 (10) | . | . |
| Grass pasture | 0.10, 0.09 (7) | . | . |
| Idle cropland | -0.03, 0.02 (5) | . | . |
| Mallard, *Anas platyrhynchos* | . | 0.85, 0.13 (58) | . |
| Song Sparrow, *Melospiza melodia* | . | -0.75, 0.12 (14) | . |
| Hairy Woodpecker, *Dryobates villosus* | . | 0.50, 0.10 (3) | . |
| Belted Kingfisher, *Megaceryle alcyon* | . | 0.48, 0.07 (2) | . |
| Carolina Wren, *Thryothorus ludovicianus* | . | 0.29, 0.02 (2) | . |
| Wood Duck, *Aix sponsa* | . | 0.06, 0.49 (2) | . |
| Tree Swallow, *Tachycineta bicolor* | . | -0.69, 0.18 (2) | . |
| White-breasted Nuthatch, *Sitta carolinensis* | . | -0.31 (1) | . |
| Northern Cardinal, *Cardinalis cardinalis* | . | -0.34 (1) | . |
| Pileated Woodpecker, *Dryocopus pileatus* | . | -0.38 (1) | . |
| American Crow, *Corvus brachyrhynchos* | . | -0.39 (1) | . |
| Eastern Towhee, *Pipilo erythrophthalmus* | . | -0.65 (1) | . |
| Canada Goose, *Branta canadensis* | . | -0.68 (1) | . |
